# A Critical Re-evaluation of the Somatic Mutation Theory (SMT) Conceptual and Quantitative Limits of the Cell-Centric Paradigm

**DOI:** 10.1101/2025.10.17.683033

**Authors:** Finn Ernst Olsen, Christian Liisberg

## Abstract

The Somatic Mutation Theory (SMT) frames cancer as a stochastic process driven by the accumulation of random mutations in somatic cells. Using a Poisson–Erlang waiting-time model, we test whether empirical mutation rates and cell-division frequencies permit such multi-hit carcinogenesis within biologically realistic timescales. Even under optimistic assumptions, expected waiting times exceed progenitor-cell lifespans by several orders of magnitude, rendering sequential multi-hit carcinogenesis statistically and biologically implausible. The analysis exposes SMT’s internal inconsistencies and supports a paradigm shift from mutation accumulation to communication breakdown as the root cause of cancer.

**Companion Paper Notice:** This preprint constitutes Part I of a coordinated two-part series on the origins of cancer.

Part I critically re-examines the Somatic Mutation Theory (SMT), demonstrating its conceptual and quantitative limits as a cell-centric paradigm.

Part II - *From Cell-Centric to System-Centric Carcinogenesis: The Repair and Capacity Adaptation (RCA) Conceptual Framework* (Olsen & Liisberg, 2025b; Zenodo 10.5281/zenodo.17495975) - presents the complementary systems-biology framework that explains carcinogenesis as failure of progenitor-cell maturation within a degraded connective-tissue microenvironment.

Together, the two papers mark a paradigm shift from mutation accumulation to communication-system failure as the root cause of cancer.

A layman’s summary of Part I is available as supplementary material for non-specialist readers.

## Introduction

### From Genetic Determinism to Systems Biology

For more than half a century, the Somatic Mutation Theory (SMT) has provided oncology with its dominant explanatory framework. It portrays carcinogenesis as the cumulative outcome of random mutations occurring in somatic (non-germline) cells over the human lifespan. These mutations, by altering key regulatory genes controlling the cell cycle, apoptosis, and differentiation, are thought to convert a normal cell into one that proliferates uncontrollably and invades surrounding tissues (Weinberg, 2013; Hanahan & Weinberg, The hallmarks of cancer, 2000). In this view, cancer is a cell-autonomous and bottom-up process: stochastic genetic events inside one cell initiates malignant transformation, and natural selection acting on clonal variation drives tumor evolution.

SMT’s conceptual clarity has profoundly shaped modern oncology. It guided the discovery of oncogenes and tumor-suppressor genes, fostered targeted therapies, and supplied a unifying language for genetic lesions across cancer types. Yet its very strengths - reductionism and focus on the isolated cell - now reveal intrinsic limits. The theory assumes that genetic change is the initiating cause of malignancy and that tissue architecture, extracellular communication, and systemic regulation play merely secondary roles. Emerging evidence challenges these assumptions. Malignant cells can revert to normal phenotypes when placed in healthy microenvironments (Bissell, 2011); cancer incidence patterns diverge from expectations based on stochastic mutation rates; and advances in developmental and systems biology show that cellular behavior is governed by multi-scale communication rather than by DNA sequence alone.

These findings motivate a rigorous re-evaluation of SMT’s internal logic and quantitative plausibility. The present paper pursues three goals:

1. Quantitative testing: to evaluate whether empirically observed mutation rates and cell-division frequencies permit multi-hit carcinogenesis within realistic lifespans of proliferative lineages.
2. Conceptual analysis: to examine the paradox that the very cells capable of accumulating mutations - stem and progenitor cells - already possess regulated proliferative capacity.
3. Epistemological clarification: to delimit the scope of SMT and identify where a systems-level framework becomes necessary.

By applying a simple Poisson-based model as a consistency test, we demonstrate that SMT’s stochastic mechanism fails to meet its own temporal and biological constraints. The analysis supports a paradigm shift from mutation accumulation to communication breakdown - from a purely genetic theory toward an integrative systems perspective in which carcinogenesis arises from the loss of coordination across molecular, cellular, and tissue levels.

## Background

### Historical and Conceptual Foundations of the Somatic Mutation Theory

#### The rise of the cell-autonomous paradigm

The Somatic Mutation Theory (SMT) emerged during the mid-twentieth century in parallel with the rise of molecular genetics and the discovery of DNA as the hereditary material. The new experimental capacity to culture, manipulate, and observe isolated cells revolutionized biomedical thinking. Disease processes that had once been interpreted in terms of tissues, organs, or systemic physiology were now redefined as disorders of individual cells. In this intellectual climate, it was natural to extend the logic of genetic determinism - so successful in explaining heredity - to the realm of cancer.

Early pioneers such as Nordling (Nordling, 1953) and P. Armitage (Armitage P, 1954) observed that cancer incidence increases exponentially with age, suggesting a multistage process requiring several independent “hits.” This statistical pattern was soon linked to the idea that these “hits” correspond to random mutations in somatic cells. The discovery of viral oncogenes in the 1960s and the identification of homologous proto-oncogenes in normal DNA during the 1970s consolidated the view that genetic mutation was the initiating cause of malignancy. By the 1980s, the discovery of tumor-suppressor genes such as *TP53* and *RB1* further reinforced the paradigm: cancer appeared to be a genetic disease of somatic evolution, driven by successive losses and gains of function within a single lineage of cells.

This historical development established a cell-autonomous paradigm of cancer (Armitage P, 1954; Noble, 2016; Nordling, 1953). In SMT, causation flows from the genome outward: mutations alter cellular circuits, producing uncontrolled proliferation, evasion of apoptosis, and loss of differentiation. The tissue and organismal environment are viewed as secondary - shaping selection pressures but not fundamentally determining the malignant state. In the words of Weinberg (Weinberg, 2013), the cancer cell is a “renegade” that has escaped the body’s control, driven by its own internal logic of growth.

#### The Unnamed Cell

Despite its widespread acceptance, SMT has never explicitly identified which specific kind of cell can undergo this supposed transformation. Textbooks typically state that “a normal cell acquires mutations that cause uncontrolled proliferation,” but most cells in the adult human body are terminally differentiated and incapable of division. Such cells - neurons, skeletal muscle fibers, and secretory epithelia - cannot re-enter the cell cycle without extraordinary intervention. The “normal cell” of SMT is therefore not generic but a very special subset: cells that already retain proliferative capacity. These include stem cells, which can self-renew and give rise to differentiated progeny, and progenitor (transit-amplifying) cells, which divide briefly before maturing into functional tissue cells. Only these cell types are mitotically competent and thus capable of accumulating mutations through replication errors.

Yet SMT rarely states this explicitly. The initiating cell of carcinogenesis remains an unnamed abstraction, simultaneously representing all cells and none. This conceptual vagueness allows SMT to appear universal while avoiding the biological precision that would constrain its claims. The problem is not merely semantic: without specifying the target cell, the theory cannot define the biological context of mutation accumulation or evaluate how tissue-level control mechanisms influence cellular fate.

#### The cell-centric limitation

By confining its causal domain to the somatic cell, SMT detaches carcinogenesis from the organism’s higher-order control hierarchy. In this framework, tissues are treated as passive environments rather than as active regulators of cellular behavior. The extracellular matrix (ECM), intercellular junctions, and biochemical signaling networks that normally coordinate differentiation are reduced to background conditions. Cancer, in turn, becomes a purely local genetic accident - an internal malfunction of one cell’s molecular circuitry.

This conceptual move was historically pragmatic: it enabled researchers to focus on experimentally tractable entities such as genes, signaling pathways, and cultured cells. However, it came at a cost. The cell-centric approach obscures the multi-scale nature of biological organization, in which cells exist as parts of communicating collectives governed by mechanical, chemical, and bioelectrical feedback. It also overlooks that proliferation, differentiation, and apoptosis are not autonomous programs but emergent outcomes of continuous negotiation between the cell and its environment (Bissell, 2011; Levin, 2021; Noble, 2016).

As a result, SMT offers a clear molecular narrative but an incomplete biological one. It explains *how* a cell might proliferate abnormally if it were already disconnected from its context - but not *why* or *how* such disconnection arises in the first place. In this sense, SMT does not so much explain the origin of cancer as presuppose it.

#### The epistemological consequence

The success of SMT has been both methodological and rhetorical. It unified cancer research under a single mechanistic framework and provided powerful molecular targets for therapy. Yet its dominance has also constrained theoretical diversity. When the cell is assumed to be the sole unit of causation, any anomaly must be explained by further subdivision - additional genes, mutations, or pathways - rather than by questioning the cell-autonomous premise itself.

Modern discoveries in developmental biology, morphogenesis, and systems physiology increasingly reveal that biological control is distributed, redundant, and context dependent. Cells are not isolated agents but participants in self-organizing networks. From this perspective, cancer may represent not merely a collection of mutated cells but a breakdown of communication within the organism’s regulatory hierarchy. Recognizing this shift does not invalidate SMT’s molecular insights; it simply reframes them within a broader systems context.

Having clarified the conceptual and biological ambiguities surrounding SMT’s “unnamed” initiating cell, the next step is to test whether the theory’s quantitative premises are internally consistent. If the accumulation of random mutations within somatic lineages truly underlies carcinogenesis, then the timing of such events must be compatible with the known rates of mutation, cell division, and lineage turnover in human tissues. The following section therefore applies a minimal probabilistic boundary test—using empirically derived parameters for stem, progenitor, and terminally differentiated cells—to determine whether the SMT mechanism is feasible even under its most favorable assumptions.

## Quantitative Boundaries of the Mutation Model

### Purpose and Scope

This section evaluates whether the Somatic Mutation Theory (SMT) is quantitatively compatible with empirical biological constraints.

SMT assumes that cancer arises when a single somatic cell lineage sequentially accumulates several independent “driver” mutations—typically five to seven—within a human lifetime (Armitage P, 1954; Nordling, 1953).

By combining observed mutation rates with realistic cell-division dynamics, we can test whether such a multistep sequence is feasible within biological time limits.

The full mathematical derivation and sensitivity analysis appear in Appendix A.

1. Mutation rate (μ). Each cell division introduces random copying errors at a rate of approximately 10^⁻⁹^–10^⁻⁸^ per base pair (Lynch, 2010; Tomasetti, 2015).
2. Effective driver target (T). Only a minute fraction of the genome (≈ 10^³^–10^⁵^ bp) can yield a cancer-relevant driver effect when altered (Weinberg, 2013; Hanahan & Weinberg, Hallmarks of cancer, 2011; Vogelstein, 2013).
3. Division frequency (δ). δ denotes the mean number of mitoses per lineage per year, linking stochastic mutation probability to real-time lineage turnover.
4. Archetypal somatic cell types.
  ○ Terminally Differentiated Cells (TDCs) are post-mitotic (δ = 0) and therefore excluded from SMT’s mutational domain (Weinberg, 2013).
  ○ Progenitor Cells complete ≈ six rapid divisions before differentiation or apoptosis (Rangel-Huerta & E, 2017). They accumulate only ≈ 5–10 random mutations—far fewer than the 5–7 specific drivers required by SMT.
  ○ **Stem Cells** are long-lived and self-renewing but divide slowly (δ ≈ 70 per year) and asymmetrically (Blanpain & Fuchs, 2014), reducing opportunities for sequential multi-hit accumulation.
5. Sequential requirement. All driver mutations must occur within the same continuous lineage before differentiation interrupts that continuity.

A discrete factorial–Poisson validation of the stochastic framework is provided in Appendix B.

### Mathematical Framework

The probability that any single division introduces a driver mutation is

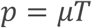

For a lineage dividing at frequency δ (divisions per year), the expected rate of driver events per lineage is

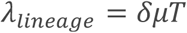

The Erlang (Gamma) expectation for *k* independent driver events within one lineage becomes

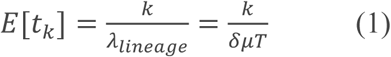

If a tissue compartment contains L parallel lineages, the first-occurrence (compartment-level) expectation is

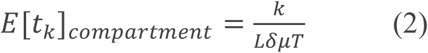

Equations 1–2 define the quantitative upper and lower bounds for stochastic multi-hit accumulation under SMT’s own assumptions.

### Quantitative Boundary Results

Even under extremely generous assumptions, SMT’s stochastic mechanism fails to meet biological time constraints (Hanahan & Weinberg, Hallmarks of cancer, 2011).

**Table 1.**
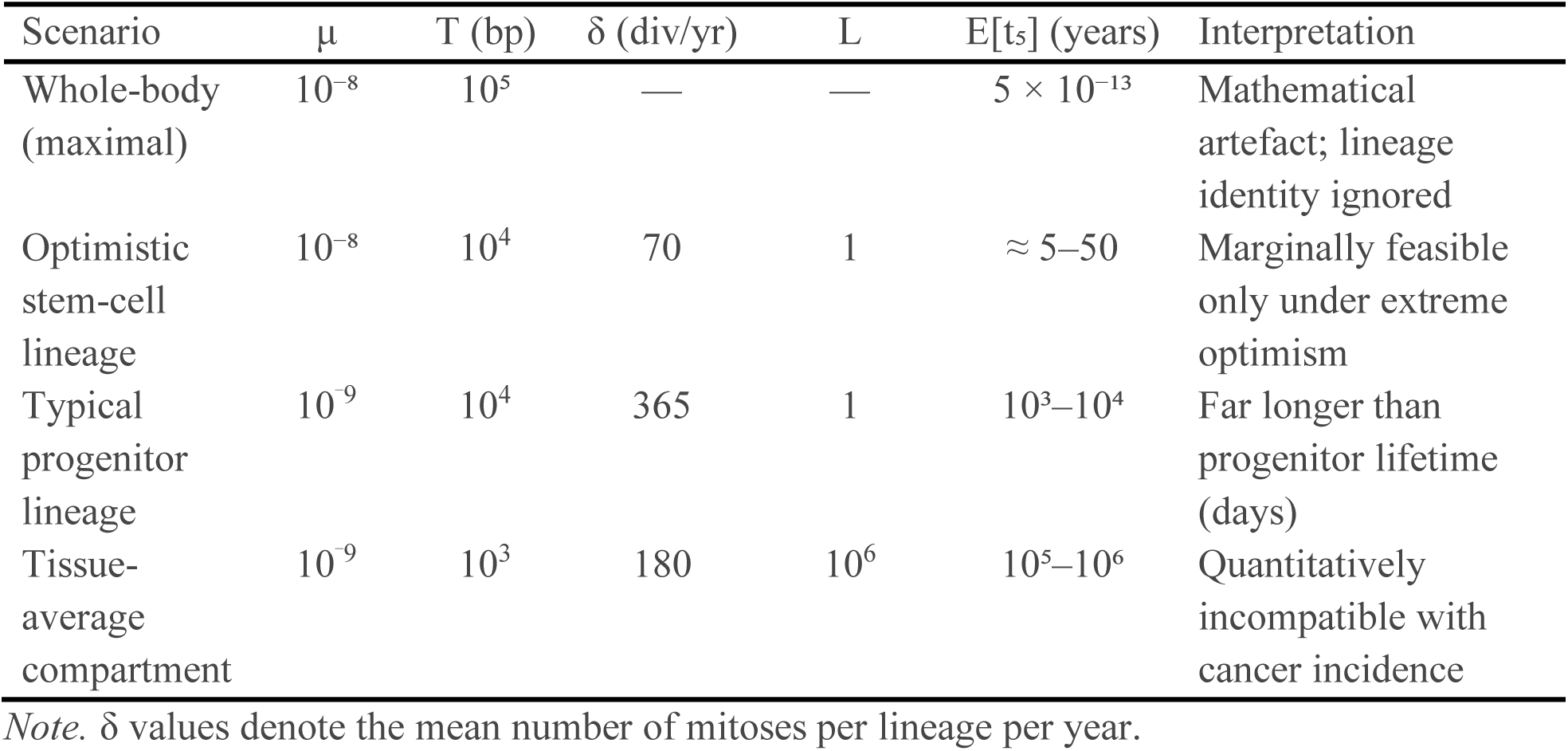
Indicative Waiting Times for Five Independent Driver Mutations.

For example, stem cells divide roughly 70 times per year, whereas progenitor cells may divide up to 365 times per year.

All waiting-time estimates are expressed in years using δ as the temporal rate parameter

### Biological Interpretation

The limiting factor is lineage lifespan, not mutation frequency.

Progenitors divide rapidly but exist too briefly; stem cells persist but divide too infrequently. No lineage provides the temporal window required for stochastic multi-hit accumulation.

Thus, the quantitative implausibility of SMT strengthens a systemic interpretation: cancer originates from disrupted differentiation control and microenvironmental signaling rather than from the random coincidence of multiple DNA errors.

*For full derivations, sensitivity tests, and biological constraints, see Appendix A*.

### The Mutation Paradox in Proliferative Archetypes

The probabilistic analysis presented above demonstrates that the accumulation of multiple independent driver mutations within a single cell lineage is exceedingly unlikely under physiological conditions. Yet even if the temporal constraints could somehow be overcome, a deeper biological paradox remains: the very cells most capable of accumulating mutations - stem and progenitor cells - already possess the ability to proliferate. Mutations that supposedly “make cells divide uncontrollably” would therefore confer a property these cells already have by design. This paradox reveals a conceptual inconsistency within the Somatic Mutation Theory (SMT) when applied to proliferative archetypes.

### Shared proliferation logic of stem and progenitor cells

Stem and progenitor cells form the regenerative backbone of all self-renewing tissues.

Stem cells are long-lived, quiescent cells capable of asymmetric division: one daughter retains stem identity while the other becomes a progenitor committed to differentiation. Progenitor cells, in turn, constitute the transit-amplifying compartment that performs a limited number of rapid divisions - typically five to seven - before maturing into functional, terminally differentiated cells.

This architecture ensures that proliferative potential is conserved but tightly controlled.

Both cell types are intrinsically competent to divide, yet neither does so autonomously. Their activity is governed by external signals from their microenvironmental niches - mechanical, biochemical, and bioelectrical cues that dictate when to divide, when to pause, and when to differentiate (Bissell, 2011; Levin, 2021). In other words, the control of proliferation resides not within the isolated genome of the cell, but within the contextual signaling network that coordinates cellular behavior across the tissue.

From this perspective, a mutation that “activates” cell division adds no new functional capacity.

At most, it could disrupt the communication that normally limits or terminates proliferation.

The problem is that this decoupling is not a single-gene event - it would require simultaneous impairment of multiple, redundant signaling pathways. SMT thus faces an internal contradiction: its proposed causal mechanism (mutation-induced uncontrolled division) overlaps with the cell’s normal proliferative function and fails to explain how contextual regulation is severed.

### The mutation paradox

The mutation paradox can be stated simply:

How can mutations in cells that already possess regulated proliferative capacity create an entirely new property of “uncontrolled proliferation”?

For a stem or progenitor cell to escape control, mutations would have to:

1. Override quiescence signals from the niche,
2. Disable differentiation pathways that terminate the cycle, and
3. Evade apoptosis triggered by improper signaling.

These requirements imply a coordinated breakdown of several independent systems - cell-cycle checkpoints (e.g., p21, p27, p57), adhesion-mediated signaling (e.g., integrins, cadherins), and differentiation regulators (e.g., Notch, Wnt/β-catenin, Hedgehog).

The probability of all these disruptions arising spontaneously in one lineage is infinitesimal, and the biological consequence would more likely be cell death or developmental arrest than malignant growth.

In essence, SMT posits that multiple destructive mutations somehow produce a new, coherent phenotype: a cell that proliferates vigorously, evades apoptosis, and maintains metabolic stability. From a systems-biology standpoint, such coordinated functionality is unlikely to emerge from random, uncorrelated genetic damage. The theory thus relies on a paradoxical assumption - that loss of regulatory information can generate higher-order functional organization.

### Weinberg’s implicit circumvention

Modern expositions of SMT, notably The Biology of Cancer (Weinberg, 2013), implicitly sidestep this paradox by redefining the effect of mutations not as the creation of new proliferative ability but as the decoupling of proliferation from environmental control.

According to this view, oncogenic mutations confer “self-sufficiency in growth signals” and “insensitivity to anti-growth signals” (Hanahan & Weinberg, Hallmarks of cancer, 2011; Bert Vogelstein, 2013)

The cancer cell is thus portrayed as autonomous - a system that no longer depends on the extracellular matrix or neighboring cells for regulatory cues.

While this formulation resolves the logical redundancy at a linguistic level, it merely relocates the problem.

The cell-autonomous model assumes that a few genetic changes can simultaneously replicate the complex, multi-directional control normally provided by the tissue microenvironment.

In practice, such decoupling would require coordinated alterations in multiple signaling axes, including growth factors, integrin-mediated anchorage, mechanical tension sensing, and polarity cues - all of which are tightly integrated in normal tissues.

Reducing this systemic regulation to a handful of gene mutations overlooks the interdependence of these networks and their emergent properties.

SMT thus explains uncontrolled growth by assuming that a cell can somehow mimic its entire environmental context through internal genetic reprogramming - a claim for which no direct experimental evidence exists.

### Empirical contradiction: microenvironmental reversion

Experimental evidence further undermines the assumption that malignant transformation is irreversible or purely cell-autonomous.

Studies by Bissell and colleagues (Bissell, 2011; Weaver, 1997) demonstrated that malignant breast epithelial cells can revert to a normal, differentiated phenotype when placed in a healthy three-dimensional extracellular matrix (ECM) environment.

Similarly, disruption of ECM integrity in otherwise normal tissue can induce invasive, disorganized growth without introducing new mutations.

These results show that contextual information from the microenvironment can override the genetic state of the cell, directing behavior through biochemical and mechanical signaling. Such findings contradict SMT’s core claim that mutations alone determine cellular phenotype. If malignant cells can be normalized by restoring environmental cues, then the primary cause of carcinogenesis must lie not in irreversible genetic damage but in the loss of regulatory communication between cells and their surroundings.

Within SMT, this reality creates a paradox: the theory attributes causation to mutations yet relies on the intact microenvironment to suppress or reveal malignant behavior.

### Summary

The mutation paradox exposes a deep logical tension at the core of the Somatic Mutation Theory.

The very cells most consistent with SMT’s probabilistic requirements - stem and progenitor cells - already possess intrinsic proliferative ability and depend on contextual signals for regulation. Mutations cannot “add” proliferation to these cells; they can only erode the communication that normally restrains it, a process far more complex than SMT’s linear, gene-centric model allows. Empirical evidence that environmental normalization can restore malignant cells to order underscores the inadequacy of cell-autonomous explanations.

The next section broadens this analysis by situating carcinogenesis within the organism’s hierarchical organization, showing how proliferation, differentiation, and repair emerge from multi-scale communication rather than from isolated genetic events.

## The Statistical and Conceptual Limits of the Somatic Mutation Theory

The Somatic Mutation Theory (SMT) provides a compelling narrative: a single somatic cell accumulates a sequence of random mutations that, over time, confer the hallmarks of malignancy. Yet when the theory is evaluated against both quantitative constraints and biological organization, its explanatory power sharply declines. The preceding sections demonstrated that the probability of acquiring the necessary combination of driver mutations within a single lineage is vanishingly small, and that the cells most compatible with SMT’s requirements - stem and progenitor cells - already possess regulated proliferative capacity. Together, these observations reveal that SMT is limited not only by statistics, but also by logic: the model presumes both an improbable sequence of events and an implausible cellular context.

This mismatch between probability, lineage identity, and observed phenotype delineates the limits of a cell-autonomous view and motivates the shift toward a systems-regulatory framework.

### The statistical boundary

The Poisson/Gamma waiting-time model clarifies the temporal horizon within which mutation-driven carcinogenesis could occur. Even under favorable assumptions - a mutation rate of 10^-8^ per base pair per division, a large mutational target (10^5^ bp), and numerous parallel lineages - the expected waiting time for five to seven driver mutations remains on the order of months to years. Under realistic physiological conditions, it extends to centuries or millennia.

These calculations underscore a simple but decisive point: the arithmetic of mutation accumulation does not match the epidemiology of cancer. Most cancers arise in mid- to late adulthood, with incidence patterns that reflect tissue renewal rates, inflammation, and aging, but not the vast combinatorial improbability that SMT implies. Even when accounting for billions of daily cell divisions across the organism, the likelihood that one lineage will independently acquire and fix the precise set of driver mutations necessary for malignancy remains infinitesimal.

This quantitative mismatch does not invalidate SMT’s individual discoveries - driver mutations undeniably exist - but it reveals that their occurrence cannot be the sole or initiating cause of cancer. The mutation rates and timescales required by SMT exceed biological reality by orders of magnitude.

### The temporal mismatch of lineage identity

Further limitation arises from the short lifespan of the relevant cell types. Progenitor cells, which form the main proliferative compartment in most tissues, undergo only a few rapid divisions before differentiating or dying. Their existence spans days, not decades. Even if the expected waiting time for multiple driver events could be shortened to a few decades, this still exceeds the lifespan of any individual progenitor lineage by several orders of magnitude.

For SMT to remain viable, it must therefore assume that successive generations of progenitors inherit partial mutational progress from an upstream, longer-lived stem cell. But this introduces another contradiction: if mutations accumulate primarily in long-lived stem cells, why do most cancers exhibit progenitor-like, not stem-like, phenotypes? SMT thus oscillates between two incompatible assumptions - mutation accumulation in stem cells and transformation in progenitor cells - without resolving how the process bridges their distinct biological roles and timescales.

### The missing role of tissue architecture and communication

The quantitative improbability and temporal mismatch both point toward a deeper conceptual omission: SMT lacks a framework for tissue-level communication and regulation.

In living organisms, cell proliferation, differentiation, and apoptosis are not autonomous properties but coordinated behaviors maintained through intercellular and extracellular signaling. The extracellular matrix (ECM), stromal fibroblasts, and mechanical and bioelectrical gradients all participate in determining whether a cell divides or differentiates (Bissell, 2011; Levin, 2021). By isolating the cell as the sole unit of causation, SMT excludes these higher-order controls from its explanatory architecture. It treats the microenvironment as a backdrop rather than a determinant. As a result, SMT cannot explain how a population of genetically normal cells can adopt malignant phenotypes under altered environmental conditions, or conversely, how genetically aberrant cells can revert to normal function when placed in a healthy context (Weaver, 1997). The theory’s gene-centric logic omits the dynamic reciprocity between cells and their surroundings that underlies tissue homeostasis.

### The logical end point of the cell-autonomous paradigm

The internal logic of SMT leads to a paradoxical conclusion: cancer is explained as a system of increasing disorder (mutation and damage) that somehow yields a new, organized, and adaptive phenotype - a self-sufficient, proliferative, invasive cell population. In thermodynamic and informational terms, this represents an inversion of biological hierarchy: random molecular errors produce coordinated, multi-trait functionality.

Such reasoning may be workable at the molecular level - individual mutations can activate signaling pathways - but it fails at the systems level, where integrated behavior emerges only through structured communication. SMT thus describes *how* genetic damage can disrupt molecular circuits but not *how* a coherent malignant phenotype arises from these disruptions. The result is a theory that is internally consistent within the cell but incomplete at the level of the organism.

### Summary

Quantitatively, SMT requires mutation rates and lineage durations incompatible with normal tissue biology. Conceptually, it assumes that a loss of regulatory information can spontaneously generate higher-order organization. The theory’s reductionist focus on the individual cell excludes the cooperative and hierarchical processes that normally govern development, repair, and differentiation.

These limitations do not diminish SMT’s historical importance but delineate its proper scope.

SMT accurately describes molecular correlates of cancer but not its emergent causation. It identifies mutations as byproducts or facilitators of malignant behavior rather than as its root cause.

Recognizing these limits does not invalidate SMT - it situates it. Cancer can no longer be explained by cell-autonomous stochastic processes alone; it must be understood as a breakdown of multi-level communication. The following sections trace how this realization has gradually forced an expansion of the mutation concept itself, as genetic determinism gives way to epigenetic and systemic regulation.

## The Expansion of the Mutation Concept

The limitations of the Somatic Mutation Theory (SMT) have not gone unnoticed within the cancer research community. Over the past two decades, as evidence has accumulated that genetic alterations alone cannot account for cancer phenotypes, the definition of “mutation” has gradually broadened. This conceptual drift reflects an effort to reconcile the persistence of SMT with findings that emphasize reversible, context-dependent, and systemic mechanisms of cellular regulation.

The result is a paradoxical expansion: the mutation concept has evolved from a precise molecular event - an irreversible change in DNA sequence - to a diffuse umbrella term encompassing epigenetic modification, chromatin remodeling, noncoding RNA dynamics, and even microenvironmental feedback. This expansion preserves SMT’s dominance linguistically but undermines its explanatory precision.

### From genetic to epigenetic alterations

Originally, SMT defined mutations as stable alterations in the nucleotide sequence of DNA - changes in the code itself.

Such mutations were assumed to be rare, permanent, and clonally inherited, providing the substrate for somatic evolution within tissues. However, research since the early 2000s has shown that heritable changes in gene expression can occur without changes to the underlying DNA sequence, through processes such as DNA methylation, histone modification, and regulation by noncoding RNAs (Feinberg A. P., 2004; Baylin, 2016).

These *epigenetic* alterations can silence tumor-suppressor genes or activate oncogenes in a manner phenotypically identical to DNA mutation - but crucially, they are reversible and context-sensitive. They can be induced by environmental stress, inflammation, nutritional state, or extracellular-matrix composition and can often be reversed by restoring normal signaling. Incorporating such mechanisms into SMT extends its reach but also changes its nature: what was once a theory of random, irreversible genetic accidents becomes a theory of regulated, adaptive information states - precisely the domain of systems biology rather than mutational stochasticity.

### Epigenetic inheritance and contextual plasticity

Epigenetic inheritance further erodes the traditional boundary between mutation and regulation. Epigenetic marks can persist across multiple cell divisions, giving the appearance of genetic heritability, yet they are dynamically maintained by enzymes whose activity depends on the cellular microenvironment. In cancer, widespread epigenetic instability - global hypomethylation alongside promoter-specific hypermethylation - has been observed across nearly all tumor types (Feinberg A. P., 1983; Sharma, 2010).

However, these patterns do not arise from random mutation; they reflect systemic stress responses and contextual reprogramming. Cells exposed to altered ECM stiffness, oxidative stress, or chronic inflammation often display reversible dedifferentiation, suggesting that what SMT describes as “mutational change” may instead represent adaptive state transitions within an unstable regulatory landscape.

By subsuming these reversible processes under the umbrella of “mutation,” SMT tacitly admits that the primary drivers of malignant behavior may not be mutational at all, but regulatory.

### Loss of falsifiability

This conceptual expansion has a critical epistemological consequence: SMT becomes non-falsifiable.

If any heritable or semi-heritable change in gene regulation can be labelled a “mutation,” the theory can no longer be disproven.

Every experimental observation - genetic, epigenetic, or environmental - can be retroactively classified as “consistent with SMT.”

The mutation concept thus evolves from a mechanistic hypothesis to a universal metaphor for all alterations in cell behavior. While this linguistic elasticity preserves the theory’s central position in oncology discourse, it dissolves its scientific rigor. A theory that can explain everything in principle explains nothing in particular.

In practice, this drift signals a tacit transition from mutation theory to regulation theory: from seeing cancer as a genomic accident to viewing it as a failure of information flow and contextual control.

### The implicit concession to systemic regulation

Ironically, by incorporating epigenetic and environmental mechanisms, SMT now approaches the very systemic perspective it once excluded.

The discovery that gene expression can be reprogrammed by extracellular cues, metabolic state, or physical context aligns more closely with tissue-level and organism-level control models than with mutation-centric causation.

In effect, the theory’s ongoing adaptation reveals its own conceptual limits: to remain viable, SMT must borrow principles from the very frameworks it was designed to replace.

This convergence does not diminish SMT’s historical importance; rather, it clarifies its position within a broader hierarchy of causation.

Mutations - whether genetic or epigenetic- can modulate cell behavior, but they act within the constraints of higher-level regulatory networks that determine cellular meaning and function. Cancer emerges not from random molecular errors alone, but from the collapse of these multi-scale communication systems.

### Summary

The gradual expansion of the mutation concept illustrates SMT’s transformation from a gene-centric to a context-inclusive paradigm.

What began as a precise hypothesis about stochastic DNA damage has evolved into a diffuse framework that encompasses reversible, environmentally induced changes in cell state.

This shift reveals that the key to understanding carcinogenesis lies not in the random accumulation of mutations, but in the failure of the regulatory architectures that ordinarily interpret and correct them.

In other words, SMT’s evolution points beyond itself: toward a systems-level view of cancer as a disorder of communication and regulation rather than of code.

The next section examines how these converging paradigms - molecular, cellular, and systemic - can be integrated into a coherent hierarchy of causation for modern oncology.

### Limitations

The present analysis evaluates SMT exclusively within a stochastic framework defined by constant mutation rate (μ), target size (T), and lineage count (L). It omits adaptive selection, clonal expansion, and chromosomal-instability bursts, all of which introduce non-random amplification and context-dependence. These simplifications ensure analytical transparency and allow a direct falsification boundary, but they necessarily restrict external predictive scope.

Consequently, the conclusions delimit rather than dismiss SMT: mutation alone provides an incomplete causal mechanism within multicellular systems.

## Discussion

### Integrating Paradigms in Modern Oncology

The Somatic Mutation Theory (SMT) has served oncology for more than half a century as both an explanatory model and a unifying research program.

Its central strength lies in its clarity: by locating causation in discrete molecular events, it has rendered the complexity of cancer experimentally tractable.

However, as shown in the preceding sections, the same reductionism that made SMT powerful also limits its scope.

Cancer does not emerge in a vacuum of molecular events; it unfolds within tissues, organs, and entire organisms that exert continuous, hierarchical control over cellular behavior.

This realization has led to the rise of complementary frameworks that restore context and communication to the center of cancer biology.

#### The domain of validity of SMT

SMT remains valid within its original domain: intracellular logic and molecular pathophysiology. Mutations in key regulatory genes - such as TP53, APC, or BRCA1 - undoubtedly contribute to the destabilization of cell-cycle control and genomic integrity.

At this level, SMT explains the mechanisms of persistence once malignancy is established.

It describes how certain clones may outcompete others, how resistance arises during therapy, and how signaling pathways adapt to selective pressure.

Within these boundaries, SMT is indispensable: it provides the molecular grammar of cancer. Yet SMT is not a theory of origins in the deeper biological sense.

It does not explain why or when a cell lineage first loses its coordination with the tissue context, nor how the multicellular architecture that normally constrains proliferation fails.

By focusing on the cell as an autonomous agent, SMT restricts itself to a single level of causation, whereas cancer is clearly a multi-level phenomenon.

#### Alternative and complementary frameworks

Several systemic theories have emerged to address this gap.

The Tissue Organization Field Theory (TOFT) proposed by Sonnenschein and Soto (1999) reframes carcinogenesis as a breakdown of tissue-level organization rather than a collection of cellular mutations.

According to TOFT, the default state of metazoan cells is proliferation, and differentiation requires active suppression by the tissue context.

Cancer thus represents a loss of that suppressive coordination rather than a gain of proliferative autonomy.

Similarly, the microenvironmental and bioelectric perspectives advanced by Bissell (Bissell, 2011) and Levin (Levin, 2021) emphasize the bidirectional flow of information between cells and their surroundings.

These frameworks portray the tissue microenvironment not as a passive scaffold but as an active regulator that transmits structural, mechanical, and electrochemical signals guiding cell fate.

When these contextual signals degrade - through aging, inflammation, fibrosis, or energetic exhaustion - cells may revert to less differentiated states, producing behaviors indistinguishable from malignancy even in the absence of new mutations.

These alternative theories do not deny the molecular events described by SMT; they embed them in a larger causal architecture.

Mutations and epigenetic modifications become secondary expressions of deeper organizational failures.

#### Levels of causation in cancer biology

To integrate these frameworks coherently, we propose a hierarchical model of causation that situates SMT within a broader systems architecture (Table 2).

**Table 2.**
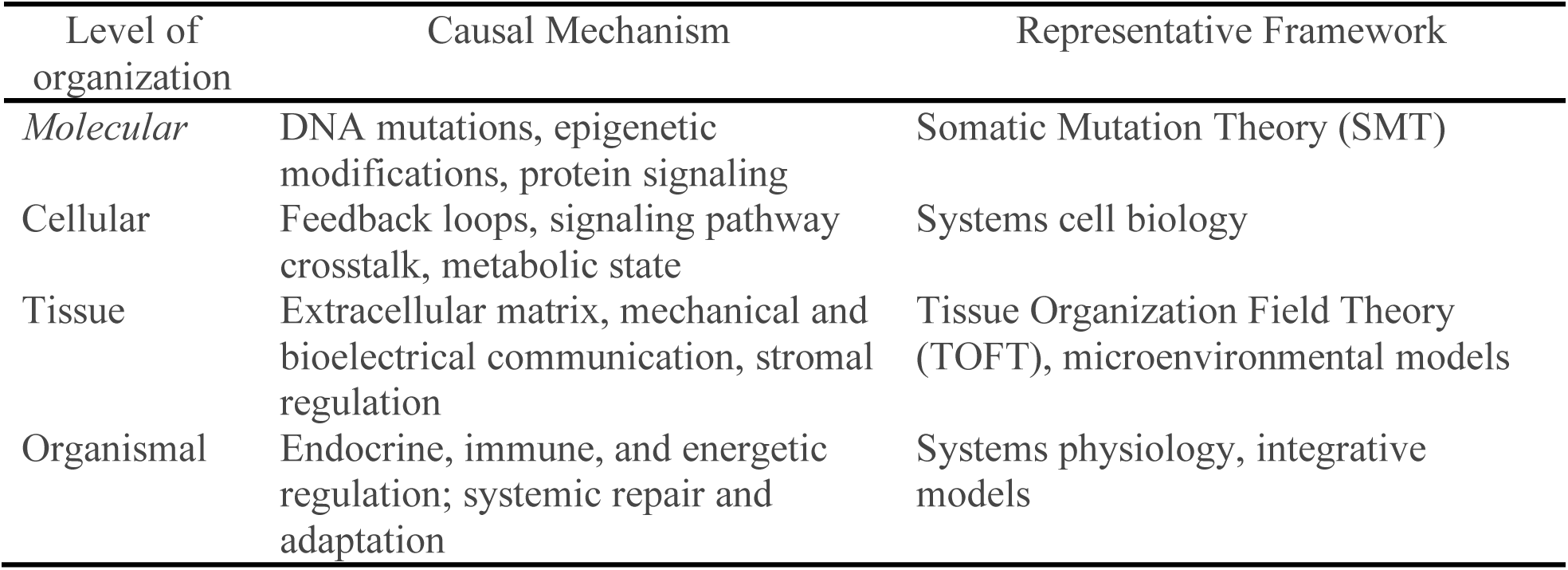
Causal Mechanism.

Causation in biology is neither strictly bottom-up nor top-down.

Instead, it is reciprocal: higher-order structures constrain and interpret molecular states, while molecular events provide the material substrate for systemic dynamics.

In this view, cancer is best understood as a failure of coordination among levels - a breakdown in the mutual communication that normally aligns cellular behavior with tissue and organismal goals.

#### Implications for research and therapy

Recognizing cancer as a multi-level disorder carries significant implications.

First, it suggests that mutation discovery alone - though essential - will never yield a complete etiology or a universal cure.

Restoring normal tissue architecture, extracellular matrix integrity, and systemic homeostasis may be as crucial as targeting genetic lesions.

Second, it highlights the importance of studying dynamic reversibility - the conditions under which malignant cells can be coaxed back toward differentiation rather than destroyed.

Finally, it invites a reorientation of cancer prevention toward the maintenance of connective-tissue health, metabolic balance, and anti-inflammatory resilience - factors that stabilize the very communication networks SMT overlooks.

#### Synthesis

SMT’s great achievement was to identify the molecular correlates of cancer; its great limitation was to mistake those correlates for ultimate causes.

The growing convergence between genetic, epigenetic, and systemic perspectives now points toward an integrative paradigm in which mutations are symptoms, not primary origins.

Cancer biology is entering a phase analogous to that faced by early genetics when molecular biology emerged: a transition from local to systemic explanations, from isolated codes to contextual meaning.

The empirical failure of stochastic sufficiency thus reframes cancer from a mutational outcome to a systems disorder of repair and capacity adaptation.

#### Summary

The Somatic Mutation Theory remains an essential component of oncology but no longer suffices as its foundation.

Future research must treat mutations, microenvironmental dynamics, and organismal regulation as co-dependent processes within a unified causal hierarchy.

The next and final section summarizes the broader implications of this re-evaluation and outlines the conceptual shift toward a systemic understanding of carcinogenesis.

## Conclusion

The Somatic Mutation Theory (SMT) transformed cancer biology.

By framing malignancy as the outcome of accumulated genetic mutations in somatic cells, it unified decades of molecular discovery into a coherent paradigm.

It explained how alterations in oncogenes and tumor suppressor genes could produce uncontrolled proliferation, resistance to apoptosis, and invasion - features long recognized as the hallmarks of cancer.

Yet, after half a century of dominance, SMT’s central assumptions have reached their empirical and conceptual limits.

Quantitatively, the stochastic accumulation of multiple driver mutations within a single lineage is statistically implausible under physiological mutation rates and cellular lifespans.

Even under optimistic conditions, the time required to acquire five or more critical mutations far exceeds the lifespan of individual progenitor cells, and far more mutations occur harmlessly than ever yield malignancy.

Biologically, SMT misidentifies the nature of proliferative control.

Stem and progenitor cells already possess the ability to divide; what constrains them is the communication network of their microenvironment.

To attribute malignant proliferation to new genetic accidents is to mistake a failure of regulation for a gain of function.

Conceptually, SMT’s reductionism confines causation to the isolated cell and excludes the higher-order organization of tissues, organs, and the organism itself.

This framework cannot explain how malignant phenotypes can be normalized in healthy environments, nor how tissue architecture and extracellular communication direct differentiation and repair.

Its recent broadening to include epigenetic, contextual, and environmental influences acknowledges this deficiency but dissolves the theory’s original clarity.

In expanding to accommodate systemic findings, SMT tacitly concedes that cancer arises from a breakdown of regulation, not solely from random mutation.

Cancer biology now stands at a crossroads.

The molecular discoveries that SMT inspired remain indispensable, but they describe only part of a larger, multi-level process.

Future progress depends on integrating genetic, epigenetic, and systemic perspectives into a single framework that recognizes the organism as an interconnected system of information exchange.

Such an approach would align oncology with modern systems biology, focusing not merely on identifying mutations but on understanding how communication, differentiation, and repair fail across scales.

A complementary systems-level framework is presented in Part II - *From Cell-Centric to System-Centric Carcinogenesis: The Repair and Capacity Adaptation (RCA) Conceptual Framework* (Olsen & Liisberg, 2025b; Zenodo 10.5281/zenodo.17495975).

The present analysis has one purpose: to clarify the internal boundaries of the Somatic Mutation Theory and demonstrate that its explanatory domain is narrower than often assumed.

Cancer cannot be reduced to a series of random genetic events; it must be understood as a loss of coordinated meaning within the living system.

### Outlook

A reorientation of oncology is underway--from mutation-centric determinism toward a systems view that integrates genetic, epigenetic, and environmental information into a coherent hierarchy of causation.

Understanding cancer as a disorder of communication opens new paths for prevention and therapy: restoring differentiation, repairing microenvironmental integrity, and maintaining energetic and structural homeostasis.

In this emerging perspective, the cell is not an isolated rebel but a component of a self-organizing organism whose health depends on the fidelity of its internal dialogue.

## Appendix A Quantitative Boundary Calculations

### A1. Core equations

Driver-mutation events are modeled as a Poisson–Erlang (Gamma) waiting-time process (Armitage P, 1954):

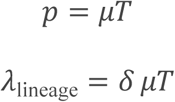

λ represents the expected driver-event rate per lineage per year

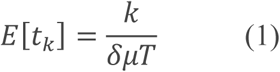

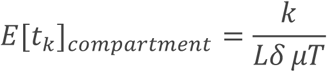

These expressions provide the expected waiting times for *k* independent driver mutations in a single lineage or across L parallel lineages.

### A2. Parameter Ranges

Model Parameters and Typical Ranges.

*Model Parameters and Typical Ranges*
| Symbol | Meaning | Typical range | Sources |
| --- | --- | --- | --- |
| $\mu$ | Per-base-pair mutation rate per division | $10^{-9} - 10^{-8}$ | (Lynch, 2010; Tomasetti, 2015) |
| $T$ | Effective driver target size (bp) | $10^3 - 10^5$ bp | (Weinberg, 2013; Vogelstein, 2013) |
| $\delta$ | Divisions per lineage per year | 70 (stem) – 365 (progenitor) | (Blanpain & Fuchs, 2014; Rangel-Huerta & E, 2017) |
| $L$ | Parallel lineages | $10^5 - 10^7$ | (Weinberg, 2013) |
| $k$ | Required driver events | 5 - 7 | (Nordling, 1953; Armitage P, 1954) |

### A3. Sensitivity and Interpretation

1. δ (lineage frequency) scales inversely with waiting time: doubling δ halves E[t₅]. Yet δ cannot rise indefinitely—biological lifespans and tissue homeostasis cap total divisions.
2. μ and T act linearly; even optimistic upper bounds shorten E[t₅] by < 10-fold, insufficient to reach observed incidence.
3. L (lineage count) affects only the *first-occurrence* boundary, not individual lineage feasibility.
4. Thus, multi-hit carcinogenesis within realistic δ, μ, T ranges remain statistically implausible.

#### A3a. Example Calculation

Using δ = 70 year⁻¹, μ = 10⁻⁸, T = 10⁴ bp, and k = 5:

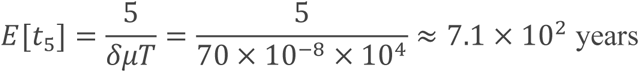

For δ = 365 year⁻¹ and μ = 10⁻⁹ (progenitor conditions):

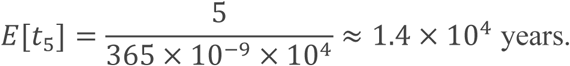

These examples illustrate the orders of magnitude shown in Table 1 of the main text.

### A4. Archetypal Cell Constraints

| Cell Type | $\delta$ (div yr<sup>-1</sup>) | Lifetime (divisions) | Feasibility for $k \geq 5$ |
| --- | --- | --- | --- |
| TDC | 0 | - | Excluded (post-mitotic) |
| Progenitor | $\approx 365$ | $\approx 6$ | Temporal window too short |
| Stem | $\approx 70$ | $> 10^4$ | Division rate too low |
| Whole-body aggregate | - | $10^{16}$ (total divisions) | Destroys lineage identity |

Only stem and progenitor compartments offer theoretical mutation opportunity, yet both fail the temporal consistency test.

### A5. Interpretive Boundary

When δ is realistic (≤ 10² year⁻¹ for stem, ≤ 4×10² year⁻¹ for progenitor), expected waiting times for five independent driver events exceed 10³–10⁶ years.

The discrepancy between these theoretical expectations and observed cancer incidence delimits SMT’s quantitative domain and redirects causation toward systemic regulation failure rather than stochastic mutation sequences.

### A6. The Clonal-Expansion Correction

A frequent objection to the quantitative boundary analysis is that once an early “driver” arises, the resulting clone proliferates faster, increasing the number of subsequent division opportunities. Formally, this would replace the constant division rate δ with a time-dependent rate δ(t) = δ₀e^{r t}.

While such feedback can reduce the expected waiting time, it introduces circular causality: the model assumes that a mutation causes self-renewal, and self-renewal is inferred from the presence of clones.

The required expansion presupposes the very phenotype—autonomous proliferation—that SMT is meant to explain.

Moreover, in intact tissues, division rates are constrained by microenvironmental control; any cell exhibiting a growth advantage is normally removed by homeostatic feedback or apoptosis.

Hence, exponential clonal amplification is biologically inconsistent with real tissue architecture. Mathematically, it rescues SMT only by abandoning its stochastic premise, shifting it toward an evolutionary or systemic framework rather than a random multi-hit mechanism.

### A7. Conclusion to Appendix A

Parameter δ bridges stochastic mutation theory to biological reality.

Even when potential corrections such as clonal expansion are considered, the multi-hit mechanism fails to achieve biologically realistic time scales without assuming prior loss of regulatory control.

Once actual division frequencies, lineage lifespans, and tissue-level feedback are imposed, SMT’s stochastic foundation collapses by several orders of magnitude.

The resulting framework delineates SMT’s quantitative boundary: beyond this point, carcinogenesis must be understood as a systemic failure of differentiation and communication rather than as the sequential accumulation of random mutations.

## Appendix B Factorial–Poisson Validation of the Stochastic Framework

### Purpose

To confirm that the quantitative boundaries derived in Appendix A are not artefacts of the continuous Poisson–Erlang approximation, a discrete factorial–Poisson formulation was examined.

This complementary analysis serves as a robustness check of the stochastic premise underlying the Somatic Mutation Theory (SMT).

### B1. Discrete combinatorial foundation

Let the genome contain *N* mutable sites (~3 × 10^⁹^ bp) and each cell division introduce *m* random point mutations (~30 per division).

The probability that any of these hits fall within the effective driver target *T* is

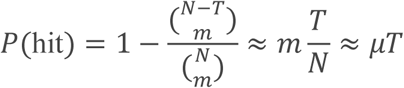

where *μ* is the per-base-pair mutation rate per division.

For a lineage requiring *k* independent driver events, the factorial–Poisson probability becomes

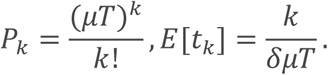

in which *δ* denotes the mean number of divisions per lineage per year.

The factorial term *k*! represents all possible permutations of the same event sequence and therefore lengthens the expected waiting time relative to the simple Poisson approximation.

### B2. Quantitative implication

Using the same parameter intervals applied in Appendix A (μ = 10^−9^ – 10^−8^; T = 10^3^ – 10^5^ bp; δ = 70 – 365 div yr^−9^), the factorial–Poisson formulation yields mean waiting times of the same order of magnitude as the Poisson–Erlang estimate.

Because *k*! grows rapidly with each additional required mutation (e.g., 5! = 120; 7! = 5040), the discrete model further enlarges the time needed for sequential multi-hit accumulation.

Thus, the factorial–Poisson treatment reinforces rather than relaxes the quantitative inconsistency between SMT’s assumptions and biological time constraints.

### B3. Interpretation

The factorial–Poisson derivation highlights the combinatorial explosion inherent in SMT’s multi-hit logic: multiplying small per-event probabilities (*μT* ≈ 10^−5^–10^−6^) across several required events (*k* = 5–7) yields astronomically low cumulative likelihoods.

Consequently, the quantitative critique presented in Appendix A is mathematically invariant to the statistical formalism chosen.

Both continuous (Poisson–Erlang) and discrete (factorial–Poisson) approaches converge on the same conclusion—that stochastic mutation accumulation within a single somatic lineage cannot plausibly account for cancer initiation within human biological timescales.

## References

Armitage P, D. R. (1954). The age distribution of cancer and a multi-stage theory of carcinogenesis. PubMed. doi:10.1038/bjc.1954.1

Baylin, S. B. (2016). Epigenetic determinants of cancer. PubMed. doi:10.1101/cshperspect.a019505

Bert Vogelstein, N. P. (2013). Cancer genome landscapes. PubMed. doi:10.1126/science.1235122

Bissell, M. &. (2011). Why don’t we get more cancer? A proposed role of the microenvironment in restraining cancer progression. PubMed. doi:10.1038/nm.2328

Blanpain, C., & Fuchs, E. (2014). Plasticity of epithelial stem cells in tissue regeneration and tumor initiation. doi:10.1038/nrm3820

Feinberg, A. P. (1983). Hypomethylation distinguishes genes of some human cancers from their normal counterparts. PubMed. doi:10.1038/301089a0

Feinberg, A. P. (2004). The history of cancer epigenetics. PubMed. doi:10.1038/nrc1279

Hanahan, A., & Weinberg, A. (2000). The hallmarks of cancer. PubMed. doi:10.1016/s0092-8674(00)81683-9

Hanahan, A., & Weinberg, A. (2011). Hallmarks of cancer. doi:10.1016/j.cell.2011.02.013

Levin, M. (2021). Bioelectric signaling: Reprogrammable circuits underlying embryogenesis, regeneration, and cancer. PubMed. doi:10.1016/j.cell.2021.02.034

Lynch, M. (2010). Rate, molecular spectrum, and consequences of human mutation. Proceedings of the National Academy of Sciences (PNAS). doi:10.1073/pnas.0912629107

Noble, D. (2016). Dance to the Tune of Life. doi:10.1017/9781316771488

Nordling, C. O. (1953). A New Theory on the Cancer-inducing Mechanism. PubMed. doi:10.1038/bjc.1953.8

Olsen, F., & Liisberg, C. (2025b). From Cell-Centric to System-Centric Carcinogenesis: The Repair and Capacity Adaptation (RCA) Conceptual Framework. Zenodo. doi:10.5281/zenodo.17495975

Rangel-Huerta, E., & E, M. (2017). Transit-Amplifying Cells in the Fast Lane from Stem Cells towards Differentiation. PubMed. doi:10.1155/2017/7602951

Sharma, S. K.–3. (2010). Epigenetics in cancer. PubMed. doi:10.1093/carcin/bgp220

Tomasetti, C. &. (2015). Cancer etiology. Variation in cancer risk among tissues can be explained by the number of stem cell divisions. PubMed. doi:10.1126/science.1260825

Vogelstein, B. &. (2013). Cancer genes and the pathways they control. Nature Medicine. doi:10.1038/nm.3167

Weaver, V. M. (1997, Jan 10). Reversion of the malignant phenotype of human breast cells in three-dimensional culture and in vivo by integrin blocking antibodies. Journal of Cell Biology. doi:10.1083/jcb.137.1.231

Weinberg, R. A. (2013). The biology of cancer (Second Edition ed.). GS Garland Science. Retrieved from https://www.amazon.com/Biology-Cancer-Robert-Weinberg/dp/0393887650

