## Supplementary material for "A Critical Re-evaluation of the Somatic Mutation Theory (SMT) Conceptual and Quantitative Limits of the Cell-Centric Paradigm": Layman version (from another version of this preprint: version 1, 2025-10-17)

### Layman Summary of “The Root Cause of Prostate Cancer: How Progenitor Cells and Microenvironment Degradation Lead to Cancer”

**Authors:** Finn Olsen & Christian Liisberg, TINAP Project
**Reference:** 251000 ver. 11.0
**Date:** October 2025

**What the paper asks**

If mutations alone cannot explain how cancer starts (as shown in our companion paper 251010), then **what actually goes wrong in the tissue?**
This paper explores whether prostate cancer could instead begin when immature repair cells - called *progenitor cells* - fail to mature properly because the surrounding connective tissue has become damaged.

---

**The key idea**

Every organ constantly repairs itself through a built-in *Repair and Capacity Adaptation (RCA)* process.

- **Stem cells** start the process.
- **Progenitor cells** do most of the work, dividing a few times and then maturing into the organ’s functional cells.
- This maturation is guided by signals from the tissue’s “scaffold,” the **extracellular matrix (ECM)** - a mesh of collagen and other proteins produced by fibroblasts.

When this micro-environment is healthy, repair is well-coordinated.
When it is degraded - by **age, inflammation, infection, obesity, alcohol, or chronic stress** - the guiding signals fade. Progenitor cells can then get **stuck halfway**, continuing to multiply instead of finishing their maturation.
These “trapped” cells behave exactly like cancer cells.

---

**Why this explains prostate cancer better**

- **Variety (heterogeneity):** different cancers reflect how far the cell got before it stalled.
- **Bone metastasis:** prostate-derived cells seek tissue environments similar to their own connective-tissue origin (both mesodermal), explaining why they home to bone.
- **Remission:** if the ECM is restored, trapped cells can mature again - showing cancer can, in principle, reverse.

---

**What this means for prevention and therapy**

Instead of only trying to kill tumour cells, treatment should also:

1. **Protect and rebuild the connective-tissue environment** (restore collagen balance, reduce stiffness, control inflammation).
2. **Support the RCA process** with energy (ATP) and essential building blocks such as amino and fatty acids.
3. **Reduce stressors** - age-related, metabolic, or psychological - that damage the ECM.

A healthy micro-environment keeps repair cells maturing normally and prevents them from turning malignant.

---

**Take-home message**

Cancer may not be a random genetic accident, but a **failure of tissue repair**.
When the prostate's repair system (RCA) loses its micro-environmental support, immature cells become trapped in growth mode.
Restoring the body's natural repair capacity - rather than focusing solely on mutation killing - may hold the key to preventing and even reversing prostate cancer.
